# High-quality, genome-wide SNP genotypic data for pedigreed germplasm of the diploid outbreeding species apple, peach, and sweet cherry through a common workflow

**DOI:** 10.1101/514281

**Authors:** Stijn Vanderzande, Nicholas P Howard, Lichun Cai, Cassia Da Silva Linge, Laima Antanaviciute, Marco CAM Bink, Johannes W Kruisselbrink, Nahla Bassil, Ksenija Gasic, Amy Iezzoni, Eric Van de Weg, Cameron Peace

## Abstract

High-quality genotypic data is a requirement for many genetic analyses. For any crop, errors in genotype calls, phasing of markers, linkage maps, pedigree records, and unnoticed variation in ploidy levels can lead to spurious marker-locus-trait associations and incorrect origin assignment of alleles to individuals. High-throughput genotyping requires automated scoring, as manual inspection of thousands of scored loci is too time-consuming. However, automated SNP scoring can result in errors that should be corrected to ensure recorded genotypic data are accurate and thereby ensure confidence in downstream genetic analyses. To enable quick identification of errors in a large genotypic data set, we have developed a comprehensive workflow. This multiple-step workflow is based on inheritance principles and on removal of markers and individuals that do not follow these principles, as demonstrated here for apple, peach, and sweet cherry. Genotypic data was obtained on pedigreed germplasm using 6-9K SNP arrays for each crop and a subset of well-performing SNPs was created using ASSIsT. Use of correct (and corrected) pedigree records readily identified violations of simple inheritance principles in the genotypic data, streamlined with FlexQTL™ software. Retained SNPs were grouped into haploblocks to increase the information content of single alleles and reduce computational power needed in downstream genetic analyses. Haploblock borders were defined by recombination locations detected in ancestral generations of cultivars and selections. Another round of inheritance-checking was conducted, for haploblock alleles (i.e., haplotypes). High-quality genotypic data sets were created using this workflow for pedigreed collections representing the U.S. breeding germplasm of apple, peach, and sweet cherry evaluated within the RosBREED project. These data sets contain 3855, 4005, and 1617 SNPs spread over 932, 103, and 196 haploblocks in apple, peach, and sweet cherry, respectively. The highly curated phased SNP and haplotype data sets, as well as the raw iScan data, of germplasm in the apple, peach, and sweet cherry Crop Reference Sets is available through the Genome Database for Rosaceae.

## Introduction

A high-quality, mostly error-free genotypic data set is imperative to obtain reliable results in many downstream genetic analyses. The results of genetic analyses can be influenced by even low rates of genotyping errors [1]. For example, the size of genetic maps and order of markers therein are affected by errors in genotypic data [2–4]. Inaccurate genotypic data will also lower the power, accuracy, and resolution of linkage studies and increase the number of false marker-locus-trait associations [5–7]. The number of observed (double) recombinants is inflated by errors in genotypic data [8]. Incorrect calling of recombinations in turn leads to incorrect determination of haploblock limits and assignment of haplotypes [9]. Finally, incorrect genotype calls can lead to incorrect imputations of missing data or even the improper adjustment of correct data to ensure the data is consistent with Mendelian inheritance [10].

There are several reasons for the occurrence of errors in a genotypic data set. Incorrect information about a sample’s identity, e.g., due to mixing up or mislabeling samples, causes an individual to be matched with the wrong data [1]. In clonally propagated crops, mislabeling errors can easily spread when individuals that are not true-to-type are used as parents or as base plants to create new propagules. Available pedigree information for an individual can be incorrect, causing incorrect enforcement of allele assignments. In fruit cultivars, numerous pedigree records have been confirmed or updated with the help of genetic markers [11–23]. Biological reasons such as unexpected mutation, insertions or deletions in the DNA sequence containing markers, and gene conversion can lead to inconsistencies in genotype calls and propagate errors through the data set [1]. Technician errors can also introduce errors in a data set, such as when lab protocols are not applied correctly (Hoffman and Amos 2005) or when multiple large data sets with disparate formats are integrated and edited. Finally, technological and software limitations and failures can also lead to the presence of errors [1].

SNPs have become the genetic marker of choice for many genetic analyses but, with their increased use and increasingly large numbers that can be generated, manual data curation has become more challenging. SNPs are ubiquitous within the genome and allow for simultaneous screening of many thousands of polymorphic loci via SNP arrays, Genotyping-By-Sequencing, or resequencing [24,25]. SNP arrays provide consistent information between individuals and have been developed for clonally propagated crops, such as the 8K apple array [26], 9K peach array [27], and 6K cherry array [28] developed by international teams led by RosBREED; the GrapeReSeq 18K Vitis array [29]; the 20K apple array developed by FruitBreedomics [30], all on the Illumina Infinium^®^ platform, and the strawberry 90K Axiom array [31], and the 480K apple array by FruitBreedomics on the Affymetrix axiom platform [32]. Genotyping each individual relies on the automated scoring of thousands of SNPs. As thousands to millions of SNPs are being assessed on a large set of individuals, even a low error rate in SNP scoring can correspond to a high absolute number of errors. As the number of SNPs on an array increase, it becomes more time-consuming and less feasible to manually review all automated SNP calls to identify potential errors.

For SNP arrays, incorrect genotype assignment using automated SNP scoring software occurs when intensity plots deviate from expected patterns. Automated genotyping is based on the association of specific alleles to different fluorescent molecules, the detection of these fluorescent molecules, the clustering of individual-marker data points according to intensity ratios between the different fluorescent dyes across multiple individuals into distinct regions of a genotype-calling space, and the final assignment of these clusters to genotypes. Examples of deviations that are observed in the intensity plots are the presence of additional clusters or clusters that have shifted from their expected location in the intensity plot. The presence of additional clusters or shifted clusters can be attributed to additional regions that bind to the SNP’s probe [33]. Sequence similarity of these regions with the intended target is caused by either local sequence repetition or presence of paralogous regions in the genome. The presence of these highly similar sequences can lead to multi-locus segregating SNP markers that cannot be adequately called. The calling of a single segregating locus might also be hampered by the background signal of targeted but non-segregating gene copies (ASSIsT Reference Manual p17 [34]). The presence of one or more additional SNPs, insertions, or deletions in the probe-binding region can lead to reduced or loss of binding affinity for the SNP’s probe and thereby to the presence of additional clusters, both of which can lead to incorrect genotype scoring of some SNPs [33].

No systematic workflow exists to efficiently detect and resolve all types of errors from a genotypic data set for pedigreed germplasm. Methods and software exist to tackle specific types of errors. For example, the ASSIsT software was developed for use with Illumina Infinium^®^ arrays to identify which SNPs show robust results, which SNPs might have genotype calling errors due to alleles with reduced affinity or null alleles, and which SNPs are monomorphic or failed completely [35]. Another example is the aggregation of linked SNPs into a single genetic locus, called haploblock, which facilitates tracking the inheritance of alleles within a pedigree and subsequent identification of inheritance inconsistencies [36]. Despite the existence these and other methods and software, an effective way to combine these methods has not been described.

Here we describe a curation workflow for high-resolution genetic marker data that identifies and resolves errors to obtain a robust set of genotypic data. The workflow maximizes the genotypic data obtained from high-throughput genome-scanning tools while minimizing the time needed to identify and remove errors. The workflow resulted from curation needs in the multi state and multi-crop USDA-SCRI project RosBREED [37–39] and the European project FruitBreedomics [40–42]. The workflow is demonstrated for three tree fruit crops, apple, peach, and sweet cherry, using the RosBREED germplasm sets [43]. The resulting genotypic data sets can be used by researchers to reconstruct pedigrees, establish quantitative genetic relationships, identify and validate quantitative trait loci (QTLs), and trace allele sources, leading to valuable practical and scientific genetic insights – with high confidence in the obtained results.

## Material and Methods

### Plant material

The apple, peach, and sweet cherry collections used in this study, referred to as the ‘Crop Reference Sets’, were created to represent U.S. breeding germplasm [43] for the RosBREED project [37] (www.rosbreed.org) and consisted of 451, 426, and 269 individuals for apple, peach, and sweet cherry, respectively (Tables S1-S3). Three apple breeding programs (Washington State University, the University of Minnesota, and Cornell University), three peach breeding programs (University of Arkansas, Clemson University, and Texas A&M University), and one sweet cherry program (Washington State University) each contributed additional germplasm to complement the Crop Reference Sets and better represent their important breeding parents [43]. These additional ‘Breeding Pedigree Sets’ consisted of 172, 139, and 167 apple individuals, 117, 289, and 143, peach individuals, and 259 sweet cherry individuals, respectively. The sweet cherry Breeding Pedigree Set was later made publicly available and became part of the sweet cherry Crop Reference Set. Genotypic data of the other Breeding Pedigree Sets were included as part of the data curation but individual identities of this private germplasm are not provided.

To reduce the trimming of pedigrees (as described under ‘Haploblock and haplotype generation’ below), the genotype calls of 18 additional apple individuals genotyped with the 20K SNP array in the FruitBreedomics project [42] or genotyped with the 8K SNP array at KU Leuven, Belgium (Table S1) were added to the data set to complete genotypic data of key ancestors.

### Initial parentage information

Initial parentage information was collected as part of the germplasm creation as described by Peace and co-workers (2014) [43]. For each breeding program, breeders provided pedigree records for their seedlings, selections, and released cultivars. Other pedigree records were based on historical records and available literature and were included for all progenitors, regardless of availability so that all progenitors terminated in founders (individuals with two unknown parents).

### DNA extraction and iScan

DNA extraction was conducted for apple, peach, and sweet cherry as described by Chagné and co-workers (2012) [26], Verde and co-workers (2012) [27], and Peace and co-workers (2012) [28], respectively. Genomic DNA from each individual was purified using the E-Z 96 Tissue DNA Kit (Omega Bio-Tek, Inc., Norcross, GA, USA). DNA was quantitated with the Quant-iT™ PicoGreen^®^ Assay (Invitrogen, Carlsbad, CA, USA), using the Victor multiplate reader (Perkin Elmer Inc., San Jose, CA, USA). DNA concentrations were adjusted to a minimum of 50 ng/µl, in 5 µl aliquots. For apple, DNA samples were run on the Illumina Infinium^®^ 8K apple SNP array [26] with iScans either at the Biotechnology Platform of the Agricultural Research Council (Pretoria, South Africa) or at the Research Technology Support Facility at Michigan State University (East Lansing, MI, USA), following the manufacturer’s protocol (Illumina Inc.). For peach and sweet cherry, DNA samples were run on the 9K peach SNP array [27] and 6K cherry SNP array [28], respectively, with an iScan at the Research Technology Support Facility at Michigan State University (East Lansing, MI, USA), following the manufacturer’s protocol (Illumina Inc.).

### Initial genetic maps

For each crop, available genetic maps were used as a framework to determine the initial order of reliable SNPs. Reliable SNPs (obtained as described under ‘Subset of reliable SNP obtainment’ below) that were not present in available genetic maps were incorporated by comparing their physical positions to those of flanking SNPs that were present in available genetic maps.

For apple, an integrated genetic map based on five full-sib families with ‘Honeycrisp’ as common parent [20] was used as a framework to help align additional SNPs on the 8K array. The relative order of SNPs in the map of Howard and co-workers (2017) [20] was adjusted to be consistent with the ‘Golden Delicious’ double haploid genome sequence v1.1 [44] whenever this did not result in false detection of double recombination for the original mapping populations. Then, SNPs that were included in the iGL map [45] but not included by Howard and co-workers (2017) [20] were aligned based on relative marker order between common markers of both maps and the ‘Golden Delicious’ double haploid genome sequence v1.1 [44]. In cases of conflict between the iGL map and the reference genome, only the iGL map was used as reference. Genetic positions of newly added SNPs were determined so that, in the new map, they had the same position relative to the position of flanking markers as these SNPs did in the iGL map. Finally, any remaining unmapped SNPs were positioned based solely on relative physical positions according to the ‘Golden Delicious’ double haploid genome sequence v1.1 [44]. When the genetic position in the iGL map was known for repositioned or newly added SNPs, their genetic position in the new map was determined so that they had the same position relative to the position of flanking markers as they did in the iGL map. When no genetic position in the iGL map was available, the genetic position was determined so that, in the new map, they had the same position relative to the position of flanking markers as they did in the physical genome. In peach, genetic positions were based on the peach physical position of peach genome v2.0 [46]. The peach physical map was scaled to an approximate genetic map by using a conversion factor where every 1 Mb corresponded to 4 cM. For sweet cherry, genetic positions were determined by aligning and integrating the physical positions using peach genome v2.0 [46] with the sweet cherry ‘Regina’ × ‘Lapins’ SNP linkage map [21,47].

### Workflow procedures

Throughout the workflow, several software packages were used. Below are described the main procedures used in the workflow, the associated software and parameter settings, and output files used. The order in which each functionality was used in the workflow is reported in Results section ‘Steps of the data curation workflow’.

#### Initial genotypic data obtainment (GenomeStudio^®^)

IScan output was converted to ‘AA’, ‘AB’, and ‘BB’ genotype calls for each SNP marker with the Genotyping module of GenomeStudio^®^ v2011.1 (Illumina Inc., San Diego, CA, USA) using a sample sheet to load sample intensities and a ‘Gen Call’ Threshold of 0.15 to assign samples to a genotype cluster. The sample sheet was adjusted in Microsoft Excel as follows before using it as input for GenomeStudio^®^:

- The sample sheet was saved as an ‘xls(x)’ file to avoid the loss of ‘SentrixBarcode’ information that occasionally occurs when saving it as a ‘.csv’ file.
- When individuals were separated over multiple iScan runs and sample sheets, the ‘[Data]’ sections of each sample sheet were combined into one.
- A copy of the ‘Sample_ID’ column in the ‘[Data]’ section was added and named ‘Sample_Original’.
- Sample names in the ‘Sample_ID’ were adjusted to remove any spaces or special characters (needed for some software) and avoid long names or names that could be interpreted as dates (or other special formats) by Excel.
- Duplicate and parental information was added to the ‘Replicate’, ‘Parent1’, and ‘Parent2’ columns considering the adjusted names in the ‘Sample_ID’ column.
- The resulting sample sheet was saved both as a ‘.xlsx’ file for future editing and as a ‘.csv’ file to serve as an input file for GenomeStudio^®^.

#### Low-quality and non-diploid sample identification (GenomeStudio^®^ and R)

Quality and ploidy were assessed using each sample’s B-allele frequencies calculated by GenomeStudio^®^. In GenomeStudio^®^, the histogram of the B-allele frequency was plotted for each individual by opening the ‘Histogram plot’ function of the ‘Full Data Table’, choosing the first individual in the ‘Columns’ section, and then choosing ‘B Allele Freq’ in the ‘Sub Columns’ section. The histogram for the ‘B-allele frequency’ could then be plotted for each individual by scrolling through the individuals in the ‘Columns’ section. Samples were considered of good quality when a clear heterozygous peak was observed around 0.5 with almost no SNPs having a B-allele frequency between 0.125 and 0.375 and between 0.625 and 0.75. In contrast, samples of poor quality showed no clear heterozygous peak around 0.5 and had many SNPs with a B allele-frequency between 0.125 and 0.375, and between 0.625 and 0.75. Individuals that showed more than three peaks in the histogram were classified as polyploid. Individuals that showed a ‘shoulder’ on the AB peak were classified as putative aneuploids and were examined further in B-allele frequency plots according to Chagné and co-workers (2015) [48], below.

To create B-allele frequency plots according to Chagné and co-workers (2015) [48], a subset of SNPs was created by applying the filter parameters described in Table S4A in the ‘SNP Table’ of GenomeStudio^®^. Next, the ‘Full Data Table’ of GenomeStudio^®^ was adjusted to only contain the B-allele frequency of each sample: in the ‘Column Chooser’ function of GenomeStudio^®^, ‘B Allele Freq’ was added to the ‘Displayed Subcolumns’ section while all other subcolumns were removed from this section. The resulting ‘Full Data Table’ was exported using the ‘export displayed data to a file’ function. The exported ‘Data Table’ was further adjusted to the following format: the first column contained the SNPs name, the second column contained the SNP’s cumulative position, and all subsequent columns contained the samples’ B-allele-frequencies.

Each SNP’s cumulative genomic position was determined as follows: the chromosome number corresponding to the SNP was multiplied by the power of ten which ensured that the outcome was larger than any possible position within any chromosome (e.g., if the largest physical position within any chromosome was 456,437 bp, all chromosome numbers were multiplied by 1,000,000 or 10^6^ as this is the first power of 10 that is larger than 456,437. Similarly, if the largest genetic position within any chromosome was 145 cM, each chromosome number was multiplied by 1000 or 10^3^). Then, the physical or genetic position within the chromosome was added to the adjusted chromosome number to obtain the cumulative genomic position of that SNP. The resulting file was then loaded into R [49].

An ad hoc R-script (Document S1) generated a pdf file that contained a plot for each individual where ‘B-allele frequency’ values were plotted for the subset of SNP markers that were ordered according to their cumulative position on a genetic linkage map or reference genome sequence. ‘B-allele frequency’ values were expected to be 0, 0.5, or 1 for diploids. Diploid samples were considered of sufficient quality when *almost no* SNPs (<0.3% of the subset) were observed between 0.125-0.375 and 0.625-0.875. In contrast, a sample was considered of intermediate or poor quality when many SNP markers (0.3%-3% and >3%, respectively) showed an intermediate or large discrepancy. For triploids, ‘B-allele frequency’ values were expected to be 0, 0.33, 0.66, and 1 for all chromosomes while values of 0, 0.25, 0.5, 0.75, and 1.0 were expected for tetraploids. Aneuploids had a diploid pattern for most chromosomes and a haploid or polyploid pattern for others. Individuals classified as poor quality, polyploid, and aneuploid were excluded from further analyses.

Samples were excluded from various input files and from the genotype clustering in GenomeStudio^®^ by choosing them in the ‘Samples Table’ and then choosing the ‘Exclude Selected Samples’. SNPs were then re-clustered by choosing the ‘Cluster All SNPs’ of the ‘Analysis’ section. All statistics were updated when prompted.

#### Subset of reliable SNP obtainment (ASSIsT)

The ‘Final report’ and ‘DNA report’ input files were created as described in the ASSIsT Reference Manual [34]. Briefly, a ‘Final Report’ and ‘DNA Report’ were generated using the ‘Report Wizard’ under the ‘Reports’ option of the ‘Analysis’ section. The best ‘redo’ was chosen based on the ‘10^th^ Percentile GC score’ and excluded samples were removed from the report. For the ‘Final Report’, ‘GTScore’, ‘Theta’, and ‘R’ were added to the default ‘Displayed Fields’ and data was grouped ‘by SNP’. For the ‘DNA Report’, samples were exported by ‘Sample ID’.

The pedigree input file was created in Excel by copying the ‘Sample_ID’, ‘Parent1’, and ‘Parent2’ columns from the ‘[Data]’ section of the sample sheet used to create the GenomeStudio^®^ project, adjusting the column names to ‘//SampleID’, ‘Mother’, and ‘Father’, respectively, and saving the resulting file as a tab-delimited text file. The (optional) map was created in Excel by having the SNP Names as given by GenomeStudio^®^ in the first column and their corresponding chromosome and position within the chromosome (either physical or genetic) as the second and third column, respectively. Column names were set to ‘//SNPid’, ‘Chromosome’, and ‘Position’ and the resulting file was saved as a tab-delimited text file

All input files were loaded into ASSIsT v1.01 [35] using the ‘Select’ button. Then, parameters were set using the ‘Set’ button as described in Table S4B depending on the ‘Population type’ used. ASSIsT distinguished eight marker classes, which were re-grouped into the following five categories:

- Robust SNPs: having less than 5% No Call Rate and all three possible clusters (AA, AB, and BB) present in the germplasm set. In ASSIsT, these SNPs were classified as ‘Robust’, ‘OneHomozygRare_HWE’, ‘OneHomozyRare_NotHWE’, and ‘DistortedAndUnexSegreg’
- Two cluster SNPs: having less than 5% No Call Rate and one of the homozygous clusters (AA or BB) absent in the germplasm set. In ASSIsT, these SNPs were classified as ‘ShiftHomo’
- Null-allele SNPs: having a probable null allele, classified as ‘NullAllele-Failed’ in ASSIsT
- Monomorphic SNPs: having no polymorphism, as in ASSIsT
- Failed SNPs: having more than 50% No Call Rate, poor clustering, or low intensity, as in ASSIsT

Results of SNP performance in ASSIsT were exported to the ‘Summary’ and ‘Custom SNP information table’. Genotype calls were saved in ‘Custom gtypes’ to be used in the R-script that checked pedigree records (described below in ‘Pedigree records verification’). PLINK input files were generated to check for unknown duplicates within the data (described below in ‘Duplicate individuals detection’) and FQ_DataPrepper input files were created to easily generate FlexQTL input files using FQDataPrepper (described below in ‘Genotyping error detection and adjustment’). Genotype calls for the ‘Robust SNPs’ category were automatically reported in ASSIsT output files whereas other categories were considered to contain failed SNPs and thus their genotype calls were not automatically reported. To include genotype calls of the ‘Two cluster SNPs’, genotype calls of such SNPs were extracted from GenomeStudio^®^ and added to the data files manually.

#### Duplicate individuals detection (GenomeStudio^®^ and Plink)

Genotypic data of known mutants and duplicates were compared to ensure their genotypic data were matching using the ‘Reproducibility and Heritability’ report of GenomeStudio^®^ (Analysis>Reports>Reproducibility and Heritability Report>with Calculating Errors). The data set was also screened for individuals with (unknown) identical genotypic data using Plink 1.9 [50] (https://www.cog-genomics.org/plink2). Plink input files generated with ASSIsT were copied into the folder that contained the PLINK executable (plink.exe). Then, a ‘command window’ or ‘PowerShell window’ was opened in this folder and the ‘plink.exe --file [*filename]* -missing-genotype - --genome full’ or ‘\plink.exe --file [*filename]* -missing-genotype - --genome full’ command was given, respectively, where [filename] was the name of the PLINK input files used. The resulting ‘plink.genome’ was opened in Excel and the ‘PI_HAT’ column was used to represent the proportion of identity-by-descent (IBD) between each pair of individuals. Pairs of individuals with an IBD proportion higher than 97% were considered to be duplicates because at this stage all known duplicates shared an IBD proportion of at least 97%. If individuals were true duplicates, only one was kept in the data set. If pedigree records differed between duplicate individuals, pedigree records were used to identify trueness-to-type as described below. True-to type individuals were kept in the data set and individuals that were not true-to-type were targeted for DNA re-sampling. Where two unselected seedlings from the same family were identified as duplicates, they were both targeted for re-sampling as it was unclear which of the two was true-to-type.

#### Pedigree records verification (GenomeStudio^®^, Cervus, and R)

Verification of pedigree records was performed by counting the Mendelian-inconsistent errors between an individual and (each of) its recorded parent(s) where genotypic data was available. These errors were genotypic data inconsistent with Mendel’s first law, i.e., alleles present in offspring but not present in either parent. First, parent-child (PC) errors between an individual and a single parent were defined as genotype calls where none of the parental alleles were present in the offspring. For example, the recorded offspring might be ‘BB’, ‘B null’, or ‘null null’ while the recorded parent was ‘AA’. In this example, neither the ‘B’ allele nor the ‘null’ alleles were present in the parent. Secondly, when both parents were known and confirmed, the combination of the two parents’ SNP data were compared to the offspring’s SNP data to identify parent-parent-child (PPC) errors. PPC errors were defined as genotype calls where at least one allele of the offspring was not present in any of its recorded parents. For example, in the case of an ‘AA’ x ‘AA’ -> ‘AB’ triplet, no PC error would be observed when checking each parent individually, as both parents could have contributed the ‘A’ allele to the offspring. However, combination of the two parents would create a PPC error as neither parent could have contributed the ‘B’ allele observed in the offspring.

Three ways to count Mendelian-inconsistent errors were compared. In GenomeStudio^®^, a ‘Reproducibility and Heritability’ (Analysis>Reports>Reproducibility and Heritability Report>with Calculating Errors) was generated to obtain the number of PC and PPC errors. Mendelian-inconsistent errors were calculated in the software Cervus [51] using default parameter settings. Third, an ad hoc R-script (Document S2) was used to check and identify PC and PPC relationships.

The ‘.gtypes’ ASSIsT output file was further adjusted to the following format: the first column contained an individual’s ‘Sample ID’, the second and third columns contained the individual’s ‘Mother ID’ and ‘Father ID’, respectively, and the subsequent columns contained the individual’s genotypic data. Any missing parental information was set to ‘-’. All alleles found in the data set were defined in the ‘AlleleList’ parameter whereas characters used for missing genotypes or missing alleles were defined in the ‘MissGT’ and ‘MissAllele’ parameters respectively. After loading all functions defined in the R-script, the ‘CheckParAll()’ function was used to identify Mendelian-inconsistent errors for individuals with at least one known parent in the data set. When an individual’s supposed parent was not genotyped but the supposed grandparents were genotyped, the grandparents-grandchild relationship was tested with the AB+AA-AA test in Excel using the template provided by van de Weg and co-workers (2018) [23].

A threshold was determined for the proportion of PC errors to confirm or reject PC relations using incompletely curated marker data. PC errors were counted for a thousand pairs of two random individuals in the data set that did not have a (known) PC relationship and for all pairs of individuals that had a known PC relationship. A separation was observed between the resulting distributions of PC errors for the two sets of individuals and a midway point between both distributions was used as threshold to reject parentage of an individual. Similarly, a threshold was determined to accept or reject the combination of two parents; observed PPC errors were counted for previously confirmed PPC relationships and a threshold set as 110% of the highest number observed PPC errors among these known relationships.

In cases of missing or erroneous parent information, efforts were made to identify the missing parent and, if not possible, to identify sets of possible grandparents. Hereto, all available selected material was examined (ancestors, direct parents, and breeding selections). In apple and peach, the ‘FindPosParComb()’ function of the ad hoc R-script (Document S2) was used to find PC and PPC relationships. The maximum number of PC errors and PPC errors to still accept a PC relationship and PPC relationship, respectively, were set with the ‘thresholdPE’ and ‘thresholdPPE’ parameters of the ‘FindPosParComb()’ function, respectively. In cherry, the software Cervus [51] was used to count these errors and determine possible parents using the default parameter settings. When no second possible parent was found in the data set, possible grandparents were identified in Excel using the template provided by van de Weg and co-workers (2018) [23]. Historic records (e.g., location and time of origin) of possible grandparents were checked to ensure feasibility. Furthermore, deduced grandparent-grandchild relationships were only kept if they did not lead to a large number of reported errors during the rest of the workflow.

Pedigree information was then updated in various input files and in GenomeStudio^®^ (Analysis>Edit Parental Relationships; then choosing individual and correct parents from drop-down menu) for further analyses. All statistics in GenomeStudio^®^ were updated when prompted.

#### Genotyping error detection and adjustment (GenomeStudio^®^, FlexQTL™, and Visual FlexQTL™)

Genotyping errors were divided in two classes: Mendelian-inconsistent errors and Mendelian-consistent errors [10]. Unlike Mendelian-inconsistent errors, Mendelian-consistent errors are errors that do not infringe upon Mendel’s first law: a child’s false allele call is present in one of the parents, but results in problematic co-segregation patterns that show unexpected double recombination between markers with successive genetic/physical positions. These double recombinations might be due to issues in ploidy, calling, marker ordering, or phasing or, occasionally, gene conversion [10] (Document S3).

For individuals with verified pedigree relationships, remaining Mendelian-inconsistent errors were detected using GenomeStudio^®^ and FlexQTL™ v0.99130. In GenomeStudio^®^, the ‘SNP Table’ was filtered for SNPs with Mendelian-inconsistent errors, the ‘Error Table’ was used to identify individuals with Mendelian-inconsistent errors, and the ‘SNP Graph’ was used to examine the reported errors. FlexQTL™ input files were prepared using FlexQTL DataPrepper v1.0.0.4 (https://www.wur.nl/en/show/FlexQTL.htm). Three input files were needed to run FlexQTL DataPrepper: a map file, a pedigree file, and a data file. The map file was obtained by adjusting the ASSIsT map input file as follows: Column names were changed to ‘MarkerId’, Group’, and ‘Position’ and the file was saved as a comma-delimited file (.csv). The pedigree file was obtained by adjusting the ASSIsT pedigree input file as follows: column names were changed to ‘Name’, ‘Parent1’, and ‘Parent2’ and the file was saved in the ‘.csv’ format. The data file was obtained by converting the ‘FlexQTLDataPrepper’ from ASSIsT to the ‘.csv’ format. The data file (.dat) generated by FlexQTL DataPrepper was adjusted to ensure all individuals had either both parents specified or none. Any individual that had only one known parent was given a dummy parent. These dummy parents, as well as any named parent not in the data set, were added to the data input file with all their genotypic data set to missing. FlexQTL™ was used to check for Mendelian-inconsistent errors (parameter settings in Table S4C). Briefly, FlexQTL™ was run through using an early stop (‘pedimapV’ parameter set to ‘2’; to stop after checking the data for inconsistencies) and allowing for segregation distortion (‘MSegDelta’ parameter set to 1). This analysis summarized for each marker and each individual how many Mendelian-inconsistent errors were observed in the ‘mconsistency.csv’ file.

Mendelian-consistent errors were detected by examining double recombinations detected over small regions (<10 cM) as reported by FlexQTL™ and Visual FlexQTL™. Parameter settings of FlexQTL™ to check for double-recombinations were the same as for Mendelian-inconsistent errors above (Table S4C). The FlexQTL™ output file named ‘DoubleRecomb.csv’ listed all singletons (single markers involved in a double recombination) in the data set. Visual FlexQTL™ instead identifies all double recombinations (including singletons) that occur within a given genetic distance. The default for this distance was 10 cM and could be changed under ‘Tools>Calculate>(Re-)Compute recombination sequences’. The report on double recombinations was created through ‘Tools>Export>Export recombination sequence file’ which provided an output file called ‘DoubleRecombinations.csv’.

Genotype calls of SNPs with Mendelian-inconsistent errors or SNPs involved in detected double recombinations were further examined in GenomeStudio^®^ using the ‘SNP Graph’. Where incorrect cluster identification was detected, clusters were manually called using the ‘SNP Graph’ and FlexQTL™ was run again to ensure errors were resolved. Individuals belonging to a single cluster were chosen using the ‘Lasso Mode’ of the ‘SNP Graph’. After ‘right-clicking’ on the ‘SNP Graph’, the ‘Define X Cluster Using Selected Samples’ was chosen where ‘X’ was the appropriate genotype cluster (‘AA’, ‘AB’, or ‘BB’). The few SNPs that could not have their genotype clusters assigned simultaneously in GenomeStudio^®^ (e.g., because clusters were too closely positioned; one of the clusters for homozygous individuals was between x=0.4 and x=0.6, which is true for part of the paralogous SNP one of the homozygous clusters according to the ASSIsT Reference Manual p14 [34]; or because null alleles were present) were genotyped as follows. Individuals belonging to a single cluster were selected using the ‘Lasso Mode’ of the ‘SNP Graph’ in GenomeStudio^®^. ‘Sample_IDs’ of the chosen individuals were transferred to Excel by highlighting the ‘Sample_ID’ column in the ‘Sample Table’, using the ‘copy’ function of the ‘Samples Table’, and pasting them into Excel. In Excel, the copied ‘SampleIDs’ were then assigned a genotype call. This process was repeated until all individuals had their genotype assigned. If genotype calls could not be accurately made, the SNP was considered to have failed and removed from the data set.

Identification of Mendelian-inconsistent and Mendelian-consistent errors were also performed at the haplotype level, conducted as described above at the single SNP level. Where an unidentified error in SNP genotype scoring was detected, the corresponding SNP genotype calls were adjusted. If the calling error occurred in a single or few individuals, haplotypes were manually adjusted to reflect the change in SNP allele. In the rare event that a large group of individuals had their SNP genotype calls adjusted, the corresponding haplotypes were re-determined using PediHaplotyper [36]. Where Mendelian-inconsistent errors were due to missing SNP alleles, the individual was compared to its parent and offspring to determine the correct haplotype. For example, if an individual had a SNP haplotype of ‘A-?-B-A’ and the haplotype was not present in either parent, but a parent had a haplotype of ‘A-A-B-A’ and no haplotype of ‘A-B-B-A’, the haplotype of the offspring would be set to ‘A-A-B-A’. If both ‘A-A-B-A’ and ‘A-B-B-A’ were present in the parent, information of flanking, linked haplotypes were checked to assess if the offspring’s haplotype could be determined by minimizing the number of recombinations. Where inconsistencies in selected material were suspected to be due to a recombination in an ungenotyped progenitor, the haploblock was split in two at the suspected recombination site to avoid tracking in downstream genetics analyses of recombination in selected material. The haplotypes for those two new haploblocks were determined again using PediHaplotyper.

#### Map error detection and adjustment (FlexQTL™, Visual FlexQTL™, and Microsoft Excel)

Where double recombinations were observed and these recombinations were not due to incorrect genotype scoring, a graphical genotyping approach was used to examine and possibly adjust SNP order in the genetic map [52]. Graphical genotyping plots were created starting from the ‘SIP_Population.csv’ output file of FlexQTL™ (Document S3). FlexQTL™ was run again to ensure the errors were resolved and only if the adjustment of the SNP order did not lead to new double recombinations, a change in order was accepted. SNPs were removed from the data set if they had unexpectedly high incidences of double recombinations that could not be resolved by repositioning the SNPs in the map. Additionally, where a SNP mapped to multiple locations in different families, the SNP was removed from the data set.

#### Haploblock and haplotype determination (FlexQTL™, Visual FlexQTL™, and PediHaplotyper)

Haploblocks were defined as regions in which no recombination was observed for selected material. For phasing, parental information in the data input file of FlexQTL™ was adjusted so that the pedigree was trimmed to remove intermediate progenitors without genotypic data unless they were represented by more than four direct offspring. Because Visual FlexQTL™ does not consider any individual without offspring (e.g., new breeding selections) in haploblock determination, dummy offspring with missing genotypic data were added for individuals that did not have any offspring in the data set yet whose recombinations were desired to contribute to determination of haploblock borders. The data was phased using FlexQTL™ (parameter settings in Table S4D). Next, Visual FlexQTL™ was used to define haploblock borders under ‘Tools>Export>Export haplotype blocks file’, creating the ‘HaploBlocks.map’ file that assigns each marker to a haploblock and could be used as input for PediHaplotyper.

For SNP phasing within haploblocks, the pedigree had to be trimmed as in haploblock determination to remove intermediate progenitors without genotypic data unless they were represented by more than four direct offspring. However, dummy offspring introduced for haploblock determination were removed again before phasing the data. FlexQTL™ was then run again (parameter settings in Table S4D), with the output file named ‘mhaplotypes.csv’, which was used as an input for PediHaplotyper.

The PediHaplotyper package [36] was loaded into R and the working directory was set to the location of the input files created above (‘HaploBlocks.map’, ‘mhaplotypes.csv’, ‘flexqtl.par’, and ‘flexqtl.sort’). In R, the function ‘fq_haplotyping_session(sessionID=’prefix”, mapfile=“HaploBlocks.map”)’ was used to create the haplotype output files in the working directory where ‘prefix’ was user-defined text that prefixed all output file names. The ‘prefix_hballeleles.dat’ output file listed the composition of each haplotype of each haploblock and the ‘prefix_flexqtl.dat’,’prefix_flexqtl.map’, and ‘prefix_flexqtl.par’ output files were used as input files for FlexQTL™ for further data curation of the haplotyped data sets (resolving both Mendelian-inconsistent and Mendelian-consistent errors as described under ‘*Genotyping error detection and adjustment’*).

#### SNP classification

A SNP classifications system was established to track clustering issues and minimize future curation of new data. SNPs that passed the filter criteria from ASSIsT and that were included in the final data set were classified into four types: type 1 SNPs had no or less than 5% call editing during the curation process and no additional genotype clusters were present; type 2 SNPs had an incorrect automated cluster identification of one of the genotype clusters (e.g., ‘AA’ cluster called as ‘AB’), showed no additional clusters, and could easily be corrected; type 3 SNPs showed additional clusters because of alleles with differential intensity signals but individuals could easily be called correctly; and type 4 SNPs had null alleles but individuals with null alleles could be distinguished easily from true homozygous individuals. Type 5 SNPs could be accurately called but their genetic or physical position could not be determined accurately and were not included in the map and final data set. Type 6 SNPs were monomorphic across all individuals. Type 7 SNPs were those considered as ‘Failed’ by ASSIsT or were removed during the workflow because their genotype calls could not be manually resolved.

### Workflow creation and implementation

A workflow was constructed by identifying necessary steps of data curation and ordering them in such a way that the amount of time needed for data curation is minimized at each step. Thus, errors addressed first were those relatively easy to identify and resolve and otherwise expected to cause problems at multiple steps. The workflow was an outcome of efforts in RosBREED and FruitBreedomics on data curation in apple, peach, and cherry. Statistics at each step of curation were determined from implementing this workflow on the RosBREED germplasm described in the ‘Plant Material’ section above.

## Results

### Steps of the data curation workflow

Initial error-detection resulted in a list of possible causes for each type of detected errors (Table 1). This list identified which issues had to be resolved first and as such resulted in the workflow described below (Figure 1, Document S3). The workflow developed had three main parts, each with multiple steps. The first main part ensures that genetic principles can be applied, the second main part applies these principles on a single marker level, and the last main part applies these principles at the haploblock level. The proposed steps within each main part are described below, as conducted for apple, peach, and sweet cherry.

**Table 1:** Errors observed during the curation process and their possible causes. Causes that should be (mostly) already resolved by the stage a researcher would start checking for specific errors are in parentheses and grey font.

| Error | Cause | Solution |
| --- | --- | --- |
| Low call rate and impossible cluster identification | Probe binding issues | Remove SNP from data set |
| Unexpected B-allele frequencies | *(Probe binding issues)* | *(Remove SNP from data set)* |
|  | Unexpected ploidy | Remove sample from data set |
|  | Low sample quality | Remove sample from data set |
| High number P(P)C errors | *(Probe binding issues)* | *(Remove SNP from data set)* |
|  | *(Low sample quality)* | *(Remove sample from data set)* |
|  | Incorrect pedigree | Adjust pedigree record |
|  | Incorrect clustering | Manually determine genotype clusters |
|  | Incorrect genotype call(s) not due to cluster issues | Adjust genotype call(s) or remove SNP from data set |
| Low number P(P)C errors | *(Probe binding issues)* | *(Remove SNP from data set)* |
|  | *(Low sample quality)* | *(Remove sample from data set)* |
|  | *(Incorrect pedigree)* | *(Adjust pedigree record)* |
|  | Incorrect clustering | Manually determine genotype clusters |
|  | Incorrect genotype call(s) not due to cluster issues | Adjust genotype call(s) |
| High number double recombinations | *(Probe binding issues)* | *(Remove SNP from data set)* |
|  | *(Low sample quality)* | *(Remove sample from data set)* |
|  | *(Incorrect pedigree)* | *(Adjust pedigree record)* |
|  | *(Unexpected ploidy)* | *(Remove sample from data set)* |
|  | Incorrect clustering | Manually determine genotype clusters |
|  | Incorrect marker position in map | Adjust marker position or remove marker if it cannot be accurately mapped |
|  | Incorrect genotype call(s) not due to cluster issues | Adjust genotype call(s) |
|  | Incorrect phasing | Find responsible individual and make genotype missing |
| Low number double recombinations | *(Probe binding issues)* | *(Remove SNP from data set)* |
|  | *(Low sample quality)* | *(Remove sample from data set)* |
|  | *(Incorrect pedigree)* | *(Adjust pedigree record)* |
|  | *(Incorrect clustering)* | *(Manually determine genotype clusters)* |
|  | **Nearby double recombination\*** | Resolve nearby double recombination |
|  | Incorrect marker position in map | Adjust marker position or remove marker if it cannot be accurately mapped |
|  | Incorrect genotype call(s) not due to cluster issues | Adjust genotype call(s) |
|  | Incorrect phasing | Wait for haploblock analysis to resolve issue |
| Incorrect haplotype determination | *(Probe binding issues)* | *(Remove SNP from data set)* |
|  | *(Low sample quality)* | *(Remove sample from data set)* |
|  | *(Incorrect pedigree)* | *(Adjust pedigree record)* |
|  | *(Incorrect clustering)* | *(Manually determine genotype clusters)* |
|  | *(Incorrect marker position in map)* | *(Adjust marker position or remove marker if it cannot be accurately mapped)* |
|  | *(Incorrect genotype call(s) not due to cluster issues)* | *(Adjust genotype call(s))* |
|  | Incorrect phasing | Manually correct phasing (determine correct haplotypes) |
|  | Recombination within haplotype | Adjust haploblock borders |

\*Nearby double recombination can occur for two adjacent markers with many double recombinations and markers with few double recombinations. However, nearby double recombinations rarely lead to a high number of double recombinations for a single marker

**Figure 1:**
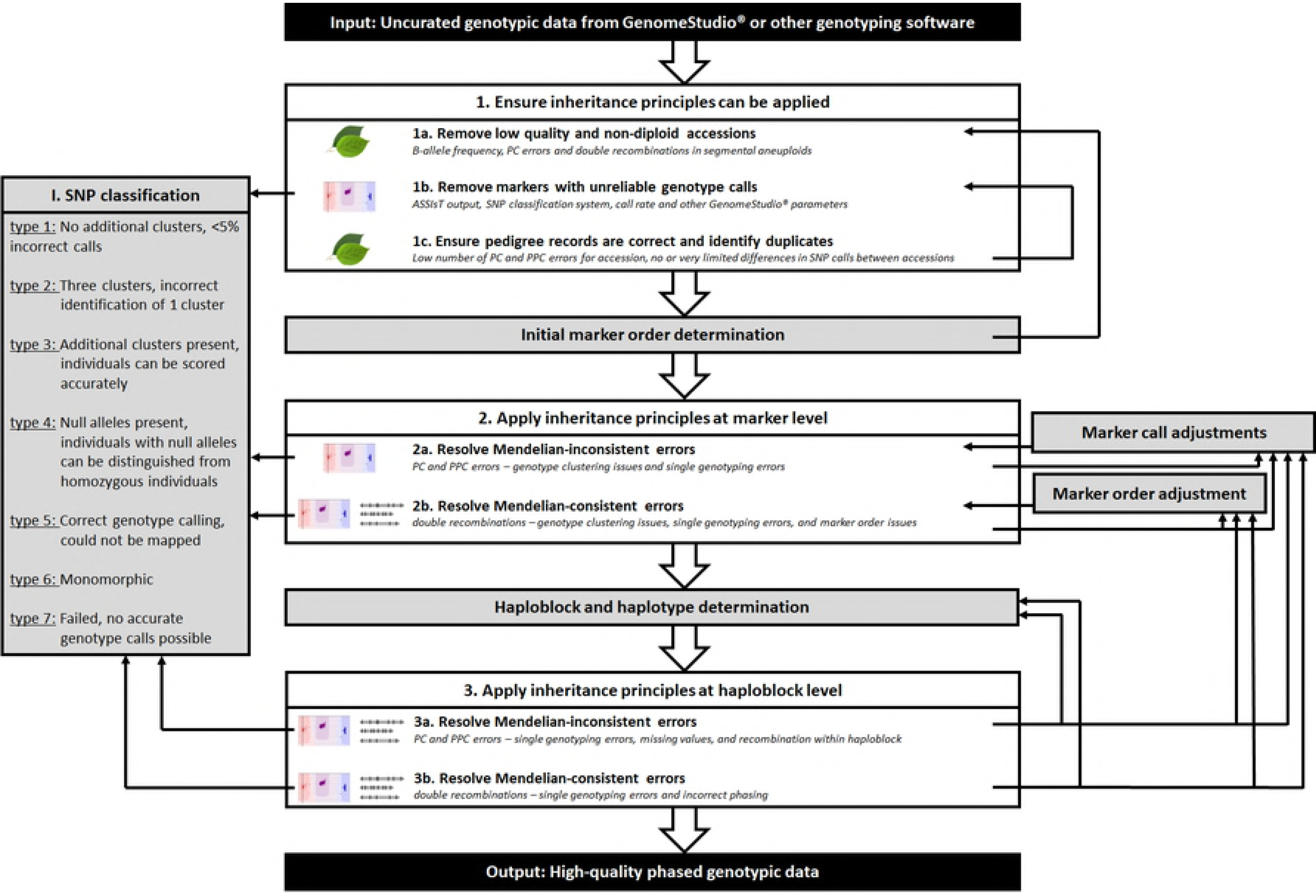
Steps of the high-resolution genotypic data curation workflow to ensure a quick and efficient curation process. Steps that identify errors are shown in white boxes; procedures needed for detecting, keeping track of, and resolving errors but do not identify errors directly are in grey boxes. After obtaining a first set of genotypic data, initial steps ensure that inheritance principles can be readily applied by removing individuals and markers that do not follow these principles and by ensuring pedigree records are correct. In the next set of steps, inheritance principles are applied at the individual marker level. In the final set of steps, these principles are applied at the haploblock level. Output used to detect and resolve observed errors at each step are given in italics. The leaf symbol indicates errors at the level of individual; the intensity plots symbol indicates errors at the level of SNP scoring; the genetic map symbol indicates errors at the level of genetically linked markers and phased alleles. When applying inheritance principles in parts 2 and 3, alleles that do not occur in an individual’s parents (‘Mendelian-inconsistent errors’) are first resolved before addressing remaining genotyping errors (‘Mendelian-consistent errors’). Several procedures, such as marker call adjustments and map order adjustments, are performed throughout the steps of the workflow to resolve errors detected. Each time after performing these common procedures, specific steps of the workflow must be repeated, forming an iterative process that ends when all errors are resolved.

#### 1. Ensuring inheritance principles can be applied

After creating an initial data set of genotypic data set in GenomeStudio^®^, a first set of analyses was performed. Because genotypic errors are identified based on principles of inheritance in diploids, individuals and markers that do not to follow these principles had to be removed first (Figure 1). When doing so, individuals with unexpected intensity patterns had to be removed first (Figure 1) as they were influencing the clustering of all individuals in the germplasm. Individuals with poor quality DNA were usually poorly genotyped, resulting in many data inconsistencies. Additionally, polyploids (individuals having one or more additional full chromosome sets) and aneuploids (individuals having an irregular number of copies for one or more chromosomes) were expected to have intensity ratios for heterozygous loci that differed from diploid individuals. Removal of individuals with poor DNA quality and suspected polyploids and aneuploids was observed to improve genotype cluster definitions and thereby the genotype calling of remaining individuals.

Once individuals with ploidy and sample quality issues were removed, a set of well performing markers had to be obtained (Figure 1). Markers with unreliable scoring were observed to lead to many inconsistencies in subsequent steps. Thus, their early removal would ensure that a relatively low number of inconsistencies remained in the data set, expected to greatly reduce the observed inconsistencies and time needed for further steps.

Identifying and correcting incorrect PC and PPC relationships was a prerequisite to using pedigree information for the identification of marker calling errors in each data set. Imposing principles of inheritance on actually unrelated individuals led to many false errors at the marker and map level. Conversely, identifying thus far unknown PC and PPC relations helped to identify errors at the marker and map level elsewhere in the data set and was expected to improve the power of downstream QTL analyses. Thus, recorded pedigree information needed to be validated and previously unknown pedigree relationships deduced before curating individual marker calls and marker order errors (Figure 1). Duplicate individuals were also detected at this stage as they could help resolve sampling errors and reduce the number of individuals needing detailed error-checking.

#### 2. Applying inheritance principles at the marker level

When Mendelian-inconsistent errors were present, at least one allele was incorrect. This issue had to be resolved before the (corrected) allele could be phased with the alleles of flanking markers. Otherwise, even the other allele, which might have been correct, could have been incorrectly phased with the alleles of flanking markers, causing additional observed but false recombinations. Thus, to minimize the time required to resolve Mendelian-consistent errors by investigating many supposed double recombinations, Mendelian-inconsistent errors had to be addressed first.

Markers with a high number of errors were investigated before markers with a relatively low number of errors among progenitors. Then, markers with a low number of errors for seedlings were investigated as they were expected to have the least effect on the remaining data set.

Any supposed double recombinations that occurred at the same region in multiple individuals had to be resolved first as they were very unlikely, could be due to a single error, and could influence a large set of individuals. Next, suspicious double recombinations that occurred over multiple loci in ancestors had to be checked, followed by singletons in ancestors. Finally, singletons in seedlings were checked, but they were expected to be the least harmful when incorrect because of little to no effect on the remaining data set.

When no genotype calling or map errors were detected, phasing errors were investigated by checking the phasing of individuals that shared the parent whose homolog was observed to have a double recombination. In the rare case that incorrect phasing by FlexQTL™ led to a double recombination in multiple individuals of a single family or parent, it was always caused by one or two individuals in which the position of (a single) recombination was incorrectly determined. In those cases, individual(s) for which the SNP was involved in a single recombination had their genotype set to missing. This adjustment led to correct phasing of all other individuals and removal of reported double recombinations. Double recombinations that were observed in a single individual and that were not due to incorrect genotype clustering or incorrect map positions were accepted as the result of true double recombination events.

#### 3. Applying inheritance principles at the haploblock level

Haploblock and haplotype determination was based on correctly identifying recombinations through correct phasing across generations and combining individual SNP alleles into haplotypes. Thus, any remaining errors at the SNP level or map level were expected to lead to errors in haploblock and haplotype determination. Therefore, all observed inconsistencies at the individual SNP level had to be resolved before inconsistencies were detected at the haploblock level. The genetic principles applied throughout the workflow are expected to also hold up at the haploblock level and therefore haplotypes had to be checked for Mendelian-consistent errors and Mendelian-inconsistent errors.

### Implementation of the workflow on RosBREED apple, peach, and sweet cherry germplasm

#### 1a. Removing samples: non-diploid individuals and low-quality samples

In apple, the ‘B allele frequency’ plot of 744 of the diploid individuals (80.7 %) was very close to that expected for diploid individuals (Figure 2A; Table S1) and results of these diploid individuals were considered to be of good quality. Another 71 individuals (7.7%) showed some variation from the expected B allele frequency, especially for homozygous SNPs, but the three genotypes could be easily distinguished (Figure 2B; Table S1) and their results quality was considered to be intermediate. Finally, 107 (11.6%) had ‘B allele frequency’ plots that showed a wide variation around the expected frequency (Figure 2C; Table S1) and their results quality was considered to be bad. No individuals with bad quality results were found for peach or sweet cherry.

**Figure 2:**
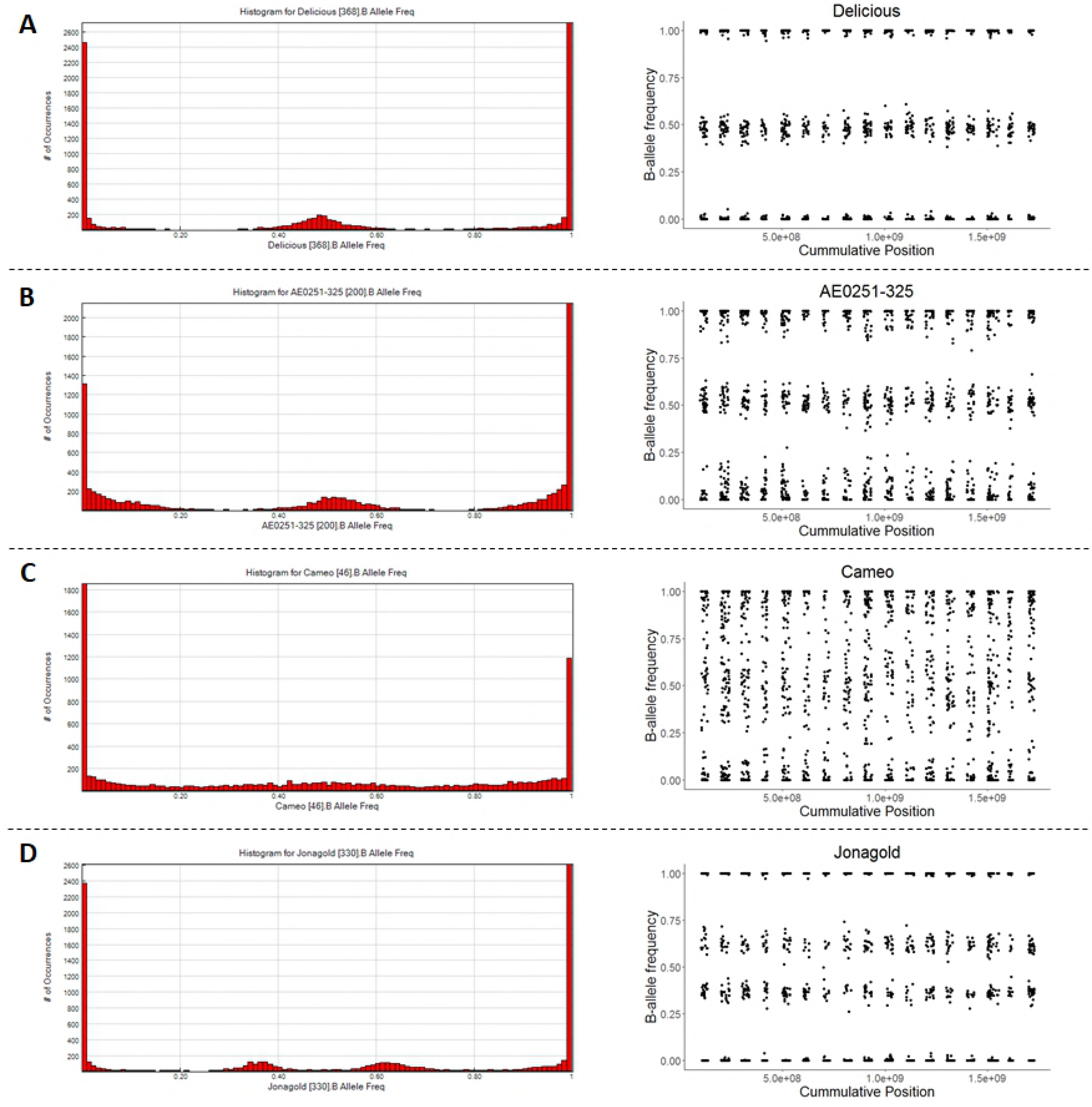
Histograms of B-allele frequency (left) and B-allele frequency for each SNP plotted against its genomic position (right). Such histograms were used to assess a sample’s genotyping quality and ploidy. Examples shown are of a sample with good quality genotype calls (panel A), with intermediate quality of genotype calls (B), with bad quality of genotype calls (C), and that is triploid (D).

For apple, most individuals with poor quality results had their DNA extracts transported outside the U.S. for genotyping and the poor results were suspected to be caused by a reduction in DNA quality due to the delay in clearing customs, while only nine individuals with poor quality were from those genotyped in the U.S. The call rate in GenomeStudio^®^ differed between the individuals that had good, intermediate, or bad quality, with the call rate dropping as the level of quality lowered (Figure S2).

For apple, five triploid individuals were identified (Table S1). One was the known triploid cultivar ‘Jonagold’ while the others were unselected seedlings (Table S1; Figure S1A). Two other unselected seedlings had their B-allele frequencies divided over 5 clusters of the GenomeStudio^®^ plot, which indicated they could be tetraploid or a mixture of two samples (Table S1; Figure S1B). No aneuploids were detected in the apple germplasm. However, one individual from the Crop Reference Set, ‘AE213-200’ and one individual of a Breeding Pedigree Set were identified as segmental aneuploids (missing one copy of a large chromosomal segment). They were undetectable in the B-allele frequency analysis and instead identified by a relatively large number of PC errors and double recombinations observed for only that chromosomal segment. No polyploids, aneuploids, or segmental aneuploids were detected in peach and sweet cherry.

The final number of individuals used in the rest of the workflow was 835, 621, and 528 for apple, peach, and sweet cherry, respectively, consisting of 139, 48, and 56 direct parents of full-sib families, ancestors, and cultivars, 76, 24, and 9 selections and 620, 548, and 463 unselected seedlings over 45, 26, and 41 families of 4–62 full-sibs, respectively (Tables S1-S3).

#### 1b. Obtaining a set of reliable SNPs

A subset of SNPs with reliable genotyping scores was obtained using ASSIsT (Table 2). Although discarded by ASSIsT, SNPs from the ‘Two cluster SNPs’ category were retained as many of them were considered to contain useful information. A total of 4636 (59%), 6098 (75%), and 1727 (30%) of the SNPs on the apple, peach, and cherry arrays, respectively, were maintained after filtering. Subsequent steps of the workflow reduced the number of SNPs in the final data set further to 3855, 4005, and 1617 for apple, peach, and sweet cherry, respectively. Thus 83%, 66%, and 91% of the SNPs retained after using ASSIsT for apple, peach, and sweet cherry, respectively, resulted in high-quality data.

**Table 2:**
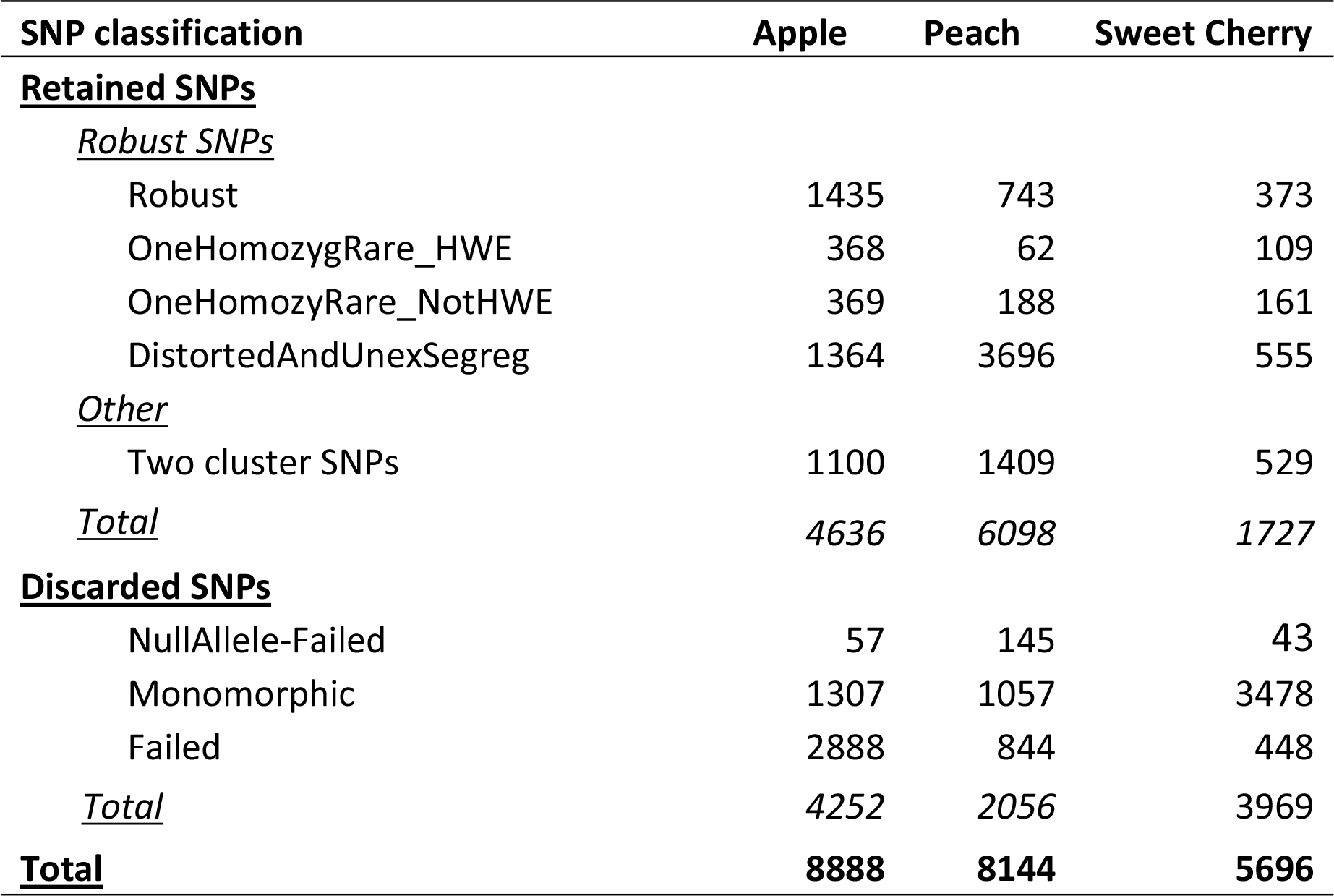
Summary of SNP classification by ASSIsT for apple, peach, and sweet cherry. SNP classifications are grouped in retained and discarded SNPs.

#### 1c. Correcting pedigree information and identifying duplicates

The number of PC errors in apple between two randomly paired individuals without PC relationship averaged 195, with a minimum of 17 (comparison between two full-sibs) and 99% of these comparisons had more than 40 errors. In contrast, average and maximum number of PC errors between two related individuals with a known PC relationship was 2 and 17, respectively, and 99% of these comparisons had less than 10 PC errors. The threshold to reject a PC relationship was set at 23 errors, which roughly corresponded to 0.5% of total markers. For 103, 66, and 22 individuals, one recorded parent was incorrect in apple, peach, and sweet cherry respectively, and for 36, 14, and zero individuals, both recorded parents were incorrect. For 106, 1, and 19 of these individuals in apple, peach, and sweet cherry, one or both of the true parent(s) was found within the germplasm set. The final number of generations spanned by the corrected pedigrees was eight, nine, and six for apple, peach and sweet cherry, respectively.

#### 2a. Finding Mendelian-inconsistent errors at the SNP level

FlexQTL™ summarized the number of Mendelian-inconsistent errors for each marker and each individual. In GenomeStudio^®^, the ‘SNP Table’ would summarize the number of P(P)C errors for each SNP and a separate ‘Error Table’ had to be consulted to determine which individuals were involved in these errors. FlexQTL™ mostly reported the error under the parent, the R-script reported the error under the offspring, and the ‘Error Table’ of GenomeStudio^®^ reported the genotypes of both parent(s) and offspring. As a consequence, errors between a single parent and multiple of its offspring would be reported as one erroneous (parental) genotype in FlexQTL™ whereas GenomeStudio^®^ reported the error for each offspring. However, FlexQTL™ did identify errors between grandparents and grandchildren when the missing parental genotype could be imputed.

FlexQTL™ detected 1209, 2230, and 686 Mendelian-inconsistent errors distributed over 541, 760, and 42 SNPs in apple, peach, and sweet cherry respectively. In apple, GenomeStudio^®^ detected 10,201 PC errors and PPC errors over 2303 SNPs. Although GenomeStudio^®^ identified which pairs of individuals led to these errors, some of the detected Mendelian-inconsistent errors did not occur in the data set due to differences in genotype scoring between ASSIsT and GenomeStudio^®^. Before removal of these Mendelian-inconsistent errors, 41,717, 29,009, and 2505 double recombinations involving a single marker were detected in FlexQTL™ in apple, peach, and sweet cherry, respectively, through the ‘DoubleRecomb.csv’ file, whereas only 6177, 4905, and 1739, respectively, of these recombinations were observed after removal of all Mendelian-inconsistent errors.

#### 2b. Identifying Mendelian-consistent errors at the SNP level

Most double recombinations that occurred in the same genomic region in many individuals could be resolved by adjusting incorrect marker calls. A total of 648, zero, and 209 markers in apple, peach, and sweet cherry, respectively, had one or more of their genotype calls adjusted to resolve double recombinations. Most other double recombinations that occurred in multiple families could be resolved by repositioning the marker in the genetic map using a graphical genotyping approach. In total, 115, zero, and zero ### SNPs were moved from their original position in the map to resolve double recombinations for apple, peach, and sweet cherry, respectively. Many recombination events that occurred in a single or few individuals over a single marker were resolved by first resolving the double recombinations that occurred in many individuals. Most of the remaining double recombinations were solved by either changing single incorrect genotype call or adjusting marker order in the map. Only a few phasing issues were observed where (almost) all offspring of a founder showed a double recombination that could be resolved by adjusting the phase of the alleles in that founder. A total of 15, 156, and 63 markers were discarded for apple, peach, and sweet cherry, respectively, because they led to unresolvable map issues. The total number of remaining reported singletons was 68, 47, 51 for apple, peach, and sweet cherry, respectively, and these were considered to be true double recombinations.

During data curation, genetic maps were generated for each crop (Tables S5-S7) by adding new SNPs to existing maps, by converting physical positions into genetic positions, and/or by updating initial genetic positions to minimize the number of double recombinations. For apple, 885 SNPs were added and 658 previously-mapped SNPs were removed as they did not perform well in our wider germplasm. Addition of SNPs at the chromosome ends enlarged the original map by 7 cM. The resulting apple map was 1179 cM long with chromosome lengths ranging from 57.6 cM (linkage group (LG) 6) to 103.6 cM (LG 15). The number of SNPs on each LG ranged from 167 SNPs on LG 6 to 359 SNPs on LG 2. The genetic map of peach was 893.2 cM long; LG 5 was the shortest (72.9 cM) and LG 1 was the longest (190.2 cM). The number of SNPs on each LG ranged from 294 on LG 5 to 772 on LG 4. In sweet cherry, chromosome lengths ranged from 56.8 cM (LG 5) to 141.2 cM (LG 1), with a total map length of 655.4 cM. The number of SNPs on each LG ranged from 137 on LG 5 to 350 on LG 1.

#### 3. Determining and resolving errors for haploblocks and haplotypes

The genetic maps of apple, peach, and sweet cherry were at first divided in 840, 103, 132 haploblocks, respectively, within which no recombination was observed in selected germplasm. After haplotype generation, 1262, 2012, and 74 Mendelian-inconsistent errors were reported by the mconsistency.csv file generated by FlexQTL™. An additional 124, 429, and 64 recombinations were detected within the haploblocks for selected germplasm, resulting in the generation of additional haploblocks. The remaining Mendelian-inconsistent errors were mostly due to missing data within a haplotype that could not be resolved automatically. This missing data within haplotypes led to the assignment of haplotype numbers that were different to parental haplotypes that were therefore perceived as errors. In addition, some inconsistencies between SNP data and haplotype data were observed after haplotype generation that were easily resolved by looking at the ‘SNP Graph’ in GenomeStudio^®^ and adjusting either the haplotype or the SNP call.

The final number of haploblocks was 964, 135, and 196 for apple, peach, and sweet cherry respectively. For apple, the genetic length of the haploblocks varied between 0 and 7.77 cM with an average of 0.3 cM, the haploblocks contained between 1 and 15 SNPs, and the haploblocks contained an average of 4 SNPs. The number of haploblocks per apple LG ranged from 42 on LG 6 to 79 on LG 15, with an average of 57 haploblocks per LG. In peach, the length of the haploblocks varied between 0 cM and 30.47 cM with an average of 5.8 cM, the haploblocks contained between 1 and 210 SNPs, and the haploblocks contained an average of 30 SNPs. The number of haploblocks per peach LG ranged from 7 on LG 5 to 37 on LG 4, with an average of 17 haploblocks per LG. For sweet cherry, haploblocks had an average length of 2.6 cM, with a minimum of 0 cM and a maximum of 15.0 cM. The average number of SNPs per sweet cherry haploblock was 8, with a minimum of 1 and a maximum of 61 SNPs. The average number of haploblocks per sweet cherry LG was 24, with a minimum of 16 haploblocks on LG 5 and LG 7 and a maximum of 47 haploblocks on LG 1.

### SNP classification system

The final number of SNPs in the haplotyped data set was 3858, 4005, and 1617 for apple, peach, and sweet cherry, respectively. A total of 3350 (87%), 4005 (100%), and 1610 (99.6%) of these SNPs were classified as type 1 SNPs, which ultimately needed editing for less than 5% of their genotype calls in apple, peach, and sweet cherry, respectively (Tables S8-10). Type 2 SNPs, for which genotype clusters were shifted, totaled 300 (8%), zero, and seven (0.4%) SNPs for apple, peach, and sweet cherry, respectively, and this shift in cluster position lead to incorrect identification of one of the three clusters in the original automatic clustering by GenomeStudio^®^. Type 3, SNPs with additional clusters, were assigned to 80 (2%), zero, and zero SNPs in apple, peach, and sweet cherry, respectively, and this presence of additional clusters led to incorrect genotype scoring of these SNPs that required subsequent curation. Type 4, SNPs with null alleles, were assigned to for 125 (3%), 145 (excluded from the final data set), and 43 (excluded from the final data set) SNPs in apple, peach, and sweet cherry, respectively, and these null alleles prevented correct automatic scoring for some individuals.

## Discussion

We established a workflow to efficiently and confidently identify and remove genotyping errors from genotyped and pedigreed germplasm sets for apple, peach, and sweet cherry. The proposed workflow (Figure 1) enables directed identification of markers and individuals with genotyping errors. It uses simple genetic principles such as inheritance of parental alleles, the co-segregation of linked markers, and the likelihood of double recombinations to find these errors. The order of steps was determined to efficiently minimize errors found in later steps and thereby minimize overall time needed to find errors in the data set. For example, in apple, any incorrect PC relationship would lead to an average of 196 reported Mendelian-inconsistent errors, and any unresolved Mendelian-inconsistent errors led to an average of 30 more reported Mendelian-consistent errors. The developed workflow was demonstrated on Illumina SNP array data and some software is specific to this platform, but the same workflow order and genetic principles are appropriate for other marker types and genotyping platforms. The workflow is especially useful when medium-and high-throughput genotyping tools are used for which checking each individual marker would be too time-consuming.

**Table 3:** Recommended software for each step of the genetic marker data curation workflow when using Illumina Infinium^®^ SNP arrays.

| Workflow step | Recommended software |
| --- | --- |
| Identify polyploids, aneuploids, and samples with low quality | GenomeStudio® to obtain B-allele frequencies,<br>R to plot B-allele frequency for each sample |
| Create subset of reliable SNPs | ASSIST |
| Identify duplicate samples | PLINK |
| Identify incorrect P(P)C relationships | GenomeStudio® |
| Identify unknown P(P)C relationships | R |
| Identify unknown grandparent-grandchild relationships | Excel* |
| Identify and resolve (remaining) Mendelian-inconsistent errors | GenomeStudio®, FlexQTL™ |
| Identify and resolve Mendelian-consistent errors | Visual FlexQTL™ + GenomeStudio® |
| Identify and correct map order inconsistencies | Visual FlexQTL™ |
| Identify phasing issues | FlexQTL™ + Visual FlexQTL™ |
| Haploblock border determination | Visual FlexQTL™ |
| Haplotype determination |  |
| - Phasing | FlexQTL™ |
| - Haplotype assignment | PediHaplotyper |
| - Curation (automated) | FlexQTL™ |

\* Template in Suppl. File 1 of Van de Weg and co-workers (2018) [23]

### Order and considerations of workflow steps

Different types of errors can be present in genotypic and pedigree data, caused by different kinds of issues (Table 1). To minimize the time needed for curation of these data, the proposed error checks need to be performed in a specific order. By first tackling issues that are common for many types of errors, subsequent curation of remaining errors becomes easier and quicker.

#### Removing individuals with low quality or irregular number of chromosome sets

The B-allele frequency plots provided a quick and easy way to identify and remove individuals with an irregular number of chromosome sets (polyploids and aneuploids) and individuals with low DNA quality. Removal of such individuals improved SNP calling and thus reduced the number of errors to be dealt with in later steps. A couple of individuals with poor quality that were originally kept, because of their importance as breeding parents, resulted in many PC errors. Making all their original SNP calls missing enabled automated imputation of most of these data points based on genetic information of relatives. Subsequent re-genotyping of these individuals matched the imputed data completely, confirming that the errors observed were due to low-quality DNA samples and not to incorrect PC relationships.

Polyploid and aneuploid individuals did not show a higher number of P(P)C errors, as expected. In contrast, these chromosome number abnormalities led to higher rates of false double recombination, either genome-wide (polyploid) or local [(segmental) aneuploids], that cannot be readily resolved other than by removal of these specific individuals.

The histogram function in GenomeStudio^®^ enabled quick identification of polyploids and individuals with very poor DNA samples without the need for additional steps in Excel, R, or other software. However, identification of aneuploids and individuals with potentially low-quality DNA samples was not as straightforward. Plotting the B-allele frequency against physical or genetic marker order (when available) required additional data manipulation and generation of the plots in software outside GenomeStudio^®^, but most of it could be automated using R and custom scripts. Therefore, we suggest using GenomeStudio^®^ for initial removal of poor-quality samples and polyploids, and afterwards, when positional information for the markers is available, screening for aneuploids with the method described by Chagné at al. (2015).

### Obtaining a set of reliable SNPs

SNPs with major scoring issues that cannot be easily resolved manually need to be removed from the data set. The early detection and removal of these unreliable SNPs greatly reduces the number of marker and map errors reported, as well as the time spent evaluating these SNPs in later workflow stages. By using ASSIsT, a quick subset of SNPs with robust genotype calls could be generated. On average across the three crops, 80% of this subset was retained in the final data set, which is lower than the 99% for single full-sib families that was reported by Di Guardo and co-workers (2015) [35]. As the number of generations and full-sib families in the germplasm increase, more SNPs with null alleles are likely to be detected and the more complicated the genotype calling of these SNPs can become. In turn, this can lead to an increased discarding of SNPs, which could explain the lower proportion of SNPs retained in our germplasm sets compared to that reported by Di Guardo and co-workers (2015) [35].

Markers with null alleles identified by ASSIsT were removed from the data set, as they could only be identified and automatically called in specific F_1_ families rather than in all families and across generations. However, many SNPs with null alleles that were later identified in the workflow could be accurately genotyped manually as long as homozygous ‘AA’ and ‘BB’ individuals could be distinguished from individuals that carried a null allele. This distinction was time-consuming and therefore we recommend saving these SNPs only when it justifies the time needed to do so. Examples when such markers can be of high value are in the construction of genetic linkage maps, even if multiple mapping populations are used [45], when they occur in a region of low coverage, or when they occur in a region of specific interest and help define additional alleles.

Very few other options exist to create a subset of high-quality genome-wide markers across pedigreed germplasm. GenomeStudio^®^ does provide several quality scores that have been used before in SNP filtering, but no guidelines exist on what threshold values to use. Using parameter thresholds regularly reported in literature [26,53–56] (GenTrain Score > 0.7, 50GC Score > 0.4, ClusterSep Score > 0.25, Call Rate > 0.9, and Minor Freq > 0.01) on the current data, the proportion of retained, unreliable, or monomorphic SNPs would be 12.3%, 23.1%, and 6.7% in apple, peach, and sweet cherry, respectively, and a large proportion of good SNPs would be discarded (27.8%, 28.2%, and 7.6%, respectively). Thus, ASSIsT greatly increased the number of reliable SNPs that were retained without reducing the quality of the subset of SNPs, making it the most efficient method to choose SNPs without prior knowledge on SNP performance.

### Updating pedigree records

As thresholds to confirm or discard historic pedigree information depends on the germplasm, genotyping platform, and data quality, they need to be assessed case-wise. A custom R-script provided quick and easy determination of the number of PC and PPC errors. However, the custom code required a significant amount of time to identify possible parents when one or both parents were unknown, especially for larger data sets. Similar issues were observed for Cervus, which took a long time to run (days) and did identify some incorrect relationships, especially for inbred material. Cervus also requires a specific data format and we experienced some problems running the software for large data sets that were not immediately resolved. GenomeStudio^®^ provided the quickest way to determine the number of PC and PPC errors, which could be determined immediately after loading the raw intensity data. However, new PC relationships could not automatically be determined and only SNPs retained by ASSIsT should be used when using GenomeStudio^®^ to determine the number of PC errors, to avoid inflating the number of PC errors. Therefore, we recommend using GenomeStudio^®^ to confirm existing pedigree records when using Illumina arrays and using an R-script to determine new, previously unknown, PC relationships. Time-consuming analyses in R could be resolved by using a subset of markers equally spread across the genome. For confirming and identifying possible grandparent-grandchild relationships, we recommend the Excel template provided by van de Weg and co-workers (2018) [23]. However, this method can misconstrue aunts-uncles/nephew-nieces and individuals with other close relationships to the target individual as grandparents. Therefore, we recommend to only use this strategy when the user has a good understanding of the germplasm such as the origin of the material and the degree of inbreeding.

Individuals with only one parent known can still be used in a pedigree-based approach to find errors in the data set, although some errors might remain unnoticed. We recommend using the ‘M_’ and ‘F_’ prefixes to the individual’s name to designate the unknown mother or father, respectively. When it is unclear whether the unknown individual is the mother or the father, the ‘UP_’ prefix can be used. Using this system instead of a non-descriptive name such as ‘dummy 1’ creates a clear connection between the individual with an unknown parent and the placeholder individual that is introduced. When the correct parent is later found, it also allows the quick replacement of the placeholder by the correct name (and corresponding genotypic data). Use of the same name for any missing parent should be avoided (e.g., using ‘dummy’ for all missing parents) unless the missing parent is unequivocally the parent of multiple individuals. If the same name is used incorrectly for multiple missing parents, the genotype of that single missing parent is expected by FlecQTLTM to be consistent with inheritance principles for all of its assigned offspring, potentially creating a large number of errors in further steps.

Although non-diploid individuals should be removed from the workflow before identifying reliable SNPs, they can have their pedigree checked if needed. Regardless of their ploidy, individuals should only contain alleles that are present in their parents. For example, a triploid individual with a marker call at one SNP of ‘AAA’ will be scored as ‘AA’, but can still not have a ‘BB’ parent. However, caution is advised as the grandparents through the parent that provided the unreduced gamete will also share a full allele set with any polyploid individual and thus these grandparents could also be incorrectly assigned as a parent of the polyploid individual. For example, the triploid ‘Zonga’ and its (diploid) grandparent ‘Cox’s Orange Pippin’ share a full allele set (through an unreduced gamete of ‘Alkmene’) and thus no PC errors are reported [57]. However, only the combination of ‘Delcorf’ and ‘Alkmene’ could explain the genotypes of the triploid ‘Zhonga’ (AB+AA-AA test [23]). Thus, for triploids, not only do parents and offspring lead to no PC errors but some grandparents do as well, and the second parent is needed to identify the true PC relationship.

### Creating or extending genetic maps

This study used available genetic maps for apple and cherry (i.e., [20,21,45,47]), integrated them when needed, and used available physical information (from [44] and [46]) to add any markers that were not already mapped. Some of these added markers were positioned at chromosome ends, which resulted in the increase of the map size by 7 cM for apple. In addition, the orientation of apple chromosome 5 was inverted here to match the orientation of the latest genome version [44]. If no genetic map is available, one will need to be constructed alongside genotypic data curation. The need for a precise genetic position of markers on the 9K peach array prompted development of consensus linkage map for peach [58] that in the future could serve as a reference map to estimate genetic positions of unmapped markers. A mapping approach for pedigreed, multi-parental maps is described by Di Pierro and co-workers (2016) [45].

### Resolving remaining Mendelian-inconsistent errors

Use of GenomeStudio^®^ for detecting Mendelian-inconsistent errors is limited to Illumina array SNPs and cannot be used for other markers or haplotypes created in later steps of the workflow. In addition, some SNPs had their SNP scoring improved with ASSIsT and manual curation, and thus the genotype scoring of GenomeStudio^®^ might not reflect the actual data. Although this latter limitation is also true when confirming pedigree data, the few differences in genotype calls between GenomeStudio^®^ and ASSIsT are not expected to alter the outcome of pedigree confirmation. In contrast, when resolving single Mendelian-inconsistent errors, it is important to know that the error is indeed present in the data set. Although Cervus counts the number of Mendelian-inconsistent errors, it does not report which markers are causing issues for which individuals, making it impractical to use to remove the remaining PC and PPCerrors. In contrast to GenomeStudio^®^, FlexQTL™ can handle multiple allele formats and is thus suited for the curation of both SNP data and haplotype data. In addition, FlexQTL™ checks for consistency over multiple generations, which enables detection of errors even if a genotype is missing in an intermediate individual. It also imputes missing data whenever possible. A disadvantage of FlexQTL™ is that it only reports one of the two individuals, often the parent, for which an error occurred; it is then up to the user to find the second individual, often the offspring, involved in the Mendelian-inconsistent error. Therefore, we recommend using FlexQTL™ to identify Mendelian-inconsistent errors and resolving them with the help of GenomeStudio^®^.

### Using map and phasing information to detect Mendelian-consistent errors

FlexQTL™ performed very accurate phasing and only a few phasing issues were noticed. Most of these phasing issues were observed as double recombinations in offspring of an individual that served as a founder. The lack of parental info for this founder provided FlexQTL™ more freedom to phase alleles, as the phasing in the founder did not need to match its parents. Incorrect phasing was most likely caused by one or very few offspring for which a true recombination occurred in the map region. In those individuals, no double recombination occurred, and the incorrect phasing inferred by FlexQTL™ minimized the interval over which the true recombination occurred. However, this minimalization of the recombination interval incorrectly specified where the recombination had occurred, causing incorrect phasing and resulting in one or multiple false double recombinations in full-and half-sibs of the individual(s) with the true recombination. Making genotype calls missing for the individual(s) with a recombination in that area enlarged the recombination interval for those individuals, but also led to correct phasing in their parent and resolved the supposed double recombinations in their full-and half-sibs. Very few other phasing issues were observed that could not be resolved on a single SNP level but were later resolved at the haploblock level. Thus, a small number of phasing issues can be accepted when moving forward to generating haploblocks and they could be nullified by FlexQTL™ by setting the parameter ‘DeleteDR’ to 1.

### Haploblock and haplotype determination

Visual FlexQTL™ showed good accuracy (between 12% and 33% of the initial haploblocks had to be divided into additional haploblocks to avoid recombination within haploblocks for selected material) in determining haploblock borders based on historic recombination events. Two reasons exist for not identifying all historic recombinations for haploblock border determination. First, Visual FlexQTL™ determines the border as the middle of the recombination interval. The more non-informative markers present in the recombination interval (due to homozygosity or lack of co-segregation (phase) information), the less likely that the middle position is the true position of the historic recombination (which determines the haploblock border). Secondly, FlexQTL™ determines haploblock borders sequentially, starting with small recombination intervals; if multiple recombinations occur in the same region, one haploblock border could suffice to account for all recombinations. This approach thus minimizes the number of recombination sites needed to explain observed segregation data. In reality, the recombinations could have occurred between different markers, requiring that region to be split in additional haploblocks to avoid recombination within haploblocks for selected material.

PediHaplotyper’s haplotypes did not always match with SNP data. In most cases, these inconsistencies were introduced during the marker consistency check with FlexQTL™ to ensure the haplotypes in an individual matched those of its parents and offspring. When the haplotype that caused the inconsistency was represented well in the pedigree, the haplotype was correct and the original genotype call for the SNP was incorrect. Thus, in these cases, haplotype curation identified additional errors in the SNP data. These errors were mostly caused by (very) small incorrectly identified genotype clusters or by single calling errors in the data set that were not detected earlier. When haplotypes in poorly represented individuals (one or two directly related individuals in the data set) showed an inconsistency with the SNP data, the SNP data was mostly correct and an error had occurred during haplotyping. The error could span multiple generations leading to inconsistencies for multiple individuals but its impact on the dataset was small as the overall representation of the incorrect haplotype was small. In the rare case that a poor representation led to incorrect haplotype determination, the actual cause of the inconsistency often remained unclear, but for some it was due to a recombination within a haploblock for an un-genotyped ancestor or one of the direct parents of such an ancestor.

Haploblock borders are not fixed and can change based on the application of the final data set and the germplasm used. For example, for QTL analyses some of the haploblocks defined here will be too large as they span multiple cM; they will show within-haploblock recombination in numerous unselected offspring thereby increasing the number of missing haplotype calls thus increasing uncertainty in QTL position (including the widening of QTL intervals). Haploblock sizes can therefore be reduced to minimize within haploblock recombination and better define QTL regions. However, when haploblocks are very small, many haploblocks will consist of only one SNP or a few SNPs, increasing data sizes (and thereby computation time in downstream analyses) and reducing the number of haplotypes per haploblock, which can reduce the suitability of the data for visual examination. Unlike the 8K apple SNP array, the 20K apple SNP array was designed to have clusters of multiple SNPs spread at approximately 1 cM intervals. A similar approach was used to create 9K add-ons for the 9K peach array and 6K cherry array [59]. This strategy supports the generation of haploblocks consisting of SNPs aggregated within 1 cM intervals while still having multiple SNPs in a single haploblock and thus multiple informative haplotypes.

Different germplasm will also lead to different haploblock borders. Currently, haploblocks are based on historic recombination events representing the U.S. breeding programs included in this study. Other breeding programs or genetic studies might have other sets of founders and thus different recombinations of relevance. Furthermore, the addition of new advanced selections and parents will introduce new recombinations in their germplasm. Finally, as the understanding of the apple, peach, and cherry germplasm increases, previously unknown progenitors, founders, and pedigree connections will be discovered, also increasing the number of observed recombinations.

Given that haploblocking is performed at a relative late stage in the workflow, haploblock borders can be altered without the need to redo all previously conducted pedigree and SNP marker curation. In fact, existing haplotype data can be converted back to phased, fully curated SNP data which, in turn, can be used to determine haplotypes for any set of haploblocks. As the SNPs are already phased and missing SNP data was imputed based on the haplotypes, haplotype determination for new haploblock borders should not create new genotyping errors in the data set. Once numbers of new recombinations are high enough to justify updating of haploblock data, part of the haploblocks and their haplotypes should be altered. PediHaplotyper supports the use of previous haplotype definitions for haploblocks that did not change in composition. Adjusted haploblocks could be marked through their names, thus providing tools to monitor new as well as previous, possibly well-known, marker alleles.

### The SNP classification system and integration of genotypic data for new germplasm into existing data sets

The established SNP classification system enables the quick creation of a subset of SNPs that require minimal or no data curation and provides a guideline on possible issues with other SNPs and how to solve them. The system should help with the quick integration of new genotypic data into existing data sets. Genotype calls for SNPs of type 1 and type 2 can be quickly integrated with high confidence in their genotype calls. Where desired, SNPs of type 3, 4, and 5 can also be integrated, but additional curation would be required. Depending on germplasm tested, these SNPs might have incorrect genotype scoring but their SNP type is an indication of why the genotype scoring is wrong and how to fix it. In other germplasm, additional SNPs in the probe or null alleles might not be present, causing SNPs that are now classified as type 4 or type 5 to give reliable results as if they were type 1 or type 2. Similarly, if germplasm is used that is unrelated to that used here, type 1 and type 2 SNPs might show additional clusters or null alleles and will require further curation. Finally, type 7 SNPs, which could not be mapped in this germplasm, might be mapped and valuable for other germplasm.

The available reference data (www.rosaceae.org), combined with the SNP classification system, will facilitate correct curation of additional genotypic data, even if the new germplasm is not directly descended. The SNP genotype calls provided here are a reference for the genotype of each observed genotype cluster in GenomeStudio^®^. In addition, SNP cluster coordinates of the latest GenomeStudio^®^ file can be imported into new projects, thus helping GenomeStudio^®^ to correctly identify clusters. Finally, the use of reference iScan data is especially useful for markers that have only two of the three clusters in a new project but all three clusters were defined in the current reference dataset. By adding reference iScan data into the new project, all three clusters will be available, ensuring correct automated genotype calling. Therefore, we recommend including available reference data when obtaining genotype calls for new germplasm.

### Data curation in apple

The need for SNP data curation in apple was increased by the whole genome duplication in the evolutionary history of apple, the relatively poorer quality of the first genome draft used for development of the 8K SNP array, and unidentified polymorphisms in the probe regions during SNP array design. The genome duplication resulted in presence of multiple highly similar sequences on different chromosomes. Indeed, a BLAST analysis against *Malus* genome v1.0 of the first 24 nucleotides of the 3’ region of arrayed SNP probes, which is most important for probe binding, showed that approximately 50% of the sequences returned multiple hits with almost all of these hits being located on multiple LGs [33]. This proportion is expected to be lower for the latest genome version [44] as most errors in assembly were removed but the proportion is expected to remain high due to chromosome and gene duplication observed in apple. Where two genomic regions are targeted by the same probe, complex cluster plots will occur if more than one of the targeted loci segregate within a single family. Such markers must be excluded from a curated data set. Multi-target markers might still be robust if they segregate at only one locus. In this case, only the cluster plot space is reduced (mostly halved), causing clusters to be located more closely to each other. In turn, this might occasionally cause separation issues. Also, some markers are lost because GenomeStudio^®^ cannot assign genotype calls for markers where one of the homozygous clusters is located at theta=0.5, the center of the x-axis, and thus these markers are considered by the software to have failed. A special case for two-locus markers occurs where each locus segregates in specific families but both loci never segregate together in the same family. In this case, genotype scoring might be performed accurately, and the SNP still needs to be present twice in the map although under different names. Two- and three-locus SNPs have been successfully mapped in the multi-family based genetic linkage map created by Di Pierro and co-workers (2016) [45]. However, in subsequent QTL mapping studies on pedigreed germplasm, such markers were excluded, as in the current study.

Several intermediate progenitors in the apple data set lacked any genotypic data and therefore the recorded link between some important breeding parents and their ancestors had to be set to unknown during haploblock and haplotype determination. For some progenitors, 20K data from the European FruitBreedomics project was available that reestablished the connection between genotyped individuals and their ancestors, but many other progenitors likely no longer exist. Individuals that were disconnected from the pedigree with little representation could not therefore have their haplotypes accurately determined using PediHaplotyper. It was, however, possible to manually determine their haplotypes based on their SNP data and haplotypes present in disconnected relatives.

### Data curation in peach

In peach, the most challenging step in the workflow was the curation of pedigree information over nine generations. Although much pedigree information is available in the literature [60], we identified incorrect parentage in the PC error analysis in cultivars and breeding selections, which we attributed to selfing or outcrossing. Incorrect pedigree records were previously reported in the UC Davis processing peach breeding program in approximately 20% of individuals, both parental and breeding selections [16]. In this work, we identified incorrect parentage in approximately 11% of the pedigree records from the three fresh market peach breeding programs, most of which were observed in breeding selections. High level of inbreeding and coancestry in the U.S cultivated peach germplasm [61] creates overlap in the ancestral generations of most U.S. peach breeding programs. Therefore, corrections in the ancestral pedigree records reported by Fresnedo-Ramírez and co-workers (2015) [16] reduced the number of errors detected here. Furthermore, intermediate parents were unavailable for genotyping, so pedigree connections were preserved by retaining pedigree information even though many intermediate progenitors were not genotyped. Finally, the presence of missing data within a haplotype resulted in Mendelian-inconsistent errors in the haploblock and haplotype generation steps, which made the haploblock data curation time-consuming.

### Data curation in sweet cherry

For the sweet cherry germplasm, the most challenging issue was the small sample size of some families (as few as four individuals), which were too small for FlexQTL™ to accurately determine linkage phase. For parents with just one genotyped offspring, phasing of the parent homologs was considered putative as recombination inherited by offspring could not be determined. For those parents with just two genotyped offspring, recombinations were arbitrarily assigned between the two offspring, as the true recombinant offspring could not be determined. In addition, scarce information on pedigrees in ancestral generations beyond about five limited further imputations in data curation, unlike for apple and peach. Various founders showed extensive regions of common haplotypes, indicating a high degree of relatedness among such founders. Some recently published haplotyping results exemplify this for the founders ‘Black Republican’ and ‘Napoleon’ [21]. Unraveling the unknown relationships among founders could thus provide useful information for future data curation in sweet cherry.

### Expectations for other crops

The proposed workflow could be applied to other diploid crops with similar breeding systems where clonally propagated relatives of current breeding material still exist. However, there are additional aspects that would need to be considered in certain circumstances that were not encountered in the present study. First, this workflow makes the assumption that there are no differences in the true SNP map order among individuals of a species. In interspecific crosses where there can be differences in chromosome arrangements between parental species, the different SNP order or indel variation among individuals could result in additional perceived double recombinations or other difficulties in following this workflow. Additionally, this workflow assumes that there is sufficient marker information to correctly identify pedigree relationships and assumes sufficient segregation information for validating marker order and identifying Mendelian-consistent errors. When using highly homozygous, inbred individuals, there might be too few segregating markers available to correctly identify marker order or find Mendelian-consistent errors through double recombinations. Also, for small germplasm sets, too few recombinations might be available to detect incorrect marker order. Finally, the prevalence of missing genotypic values should be sufficiently low across individuals. Unlike the SNP arrays used in this study, some genotyping methods such as Genotyping-by-Sequencing do not consistently target specific loci. This non-specificity can increase the flexibility of their use, but also raises new issues for which the current workflow would have to be adapted, including the potential decrease in accuracies of genotyping and haploblock determination due to unbalanced representation of genotyped loci, high levels of missing data, and sequencing errors.

### High-quality archived SNP and haplotype data sets

The presented genome-wide genotypic data sets for apple, peach, and sweet cherry are of very high quality, are composed of genetically complex germplasm, and contain no errors that could be determined based on pedigree information. This high quality provides confidence in the results of downstream analyses. Such confidence is important as many of these results are expected to lead to fundamental discoveries and practical breeding application. The iScan data, phased SNP, and haplotype datasets of individuals in the apple, peach, and sweet cherry crop reference sets are available through the Genome Database for Rosaceae (www.rosaceae.org).

Marker and pedigree data from germplasm subsets of the current U.S. RosBREED project, the EU-FruitBreedomics project, and other research projects have previously been curated by a precursor to the current workflow and used for the creation of a multi-family based genetic linkage maps [20,45] and in multifamily based QTL studies in apple [62–65], peach [22,66], and sweet cherry [14]. Also, elements of the workflow were used for allo-octoploid strawberry to curate Axiom-based SNP markers [31] and pedigree data that were subsequently used in multi-family based QTL analyses [67–69]. While providing high-quality data for each analysis separately, these earlier steps in data curation have helped guide and streamline the data curation workflow presented here. The current workflow and resulting data sets ensure that the same curation steps have been used across the data sets of multiple crops and that the data sets are of the same high quality.

## Conclusion

A curation workflow for genotypic data of pedigreed germplasm was generated by determining the optimal order of resolving issues and by providing a step-by-step guideline. Using simple genetic principles, errors can be found and curated in a directed and efficient way, reducing the time needed to obtain a high-quality genotypic data set. The workflow was used to obtain a SNP data set for large germplasm sets for each of apple, peach, and sweet cherry representing U.S. breeding programs based on the apple 8K SNP array, peach 9K SNP array, and cherry 6K SNP array, respectively, whose SNP data is available through this paper (www.rosaceae.org), as well as used on apple and peach germplasm sets representing European breeding programs based on the apple 20K and peach 9K arrays, whose SNP data are still private. These high-quality data sets contain the largest sets of SNPs obtained through their respective SNP arrays and will provide the foundation for confident subsequent analyses in genetic research.

## Acknowledgements

The authors want to thank Jasper Rees (Agricultural Research Council, South Africa) for his help in genotyping apple samples. This work was funded by the USDA-NIFA-Specialty Crop Research Initiative projects, RosBREED: Enabling marker-assisted breeding in Rosaceae (2009-51181-05808), RosBREED 2: Combining disease resistance with horticultural quality in new rosaceous cultivars (2014-51181-22378), USDA NIFA Hatch projects 0211277and 1014919, and the FruitBreedomics project No 265582: Integrated approach for increasing breeding efficiency in fruit tree crops (www.FruitBreedomics.com) that was co-funded by the EU seventh Framework Programme.

## Supplementary information

**Table S1**: Apple germplasm genotyped and used for data curation workflow. Individuals are split over the publicly available RosBREED Crop Reference Set, three privately held RosBREED Breeding Pedigree Sets, and genotypic data received from either KULeuven (Belgium) or the FruitBreedomics project. Except for the Breeding Pedigree sets, curated pedigree information is given for each individual. For each individual, the type of material (selected vs. unselected), the location of sampling, quality of the results, and inferred ploidy of the sample are given. For unselected seedlings, the family to which they belong is also given. For the Breeding Pedigree Sets, this information is summarized per full-sib family. If tissue was collected at the USDA germplasm repository in Geneva, a GRIN accession number is also provided. Parents highlighted in yellow did not have genotypic data and their pedigree-relationships could not be tested.

**Table S2**: Peach germplasm genotyped and used for curation workflow. Individuals are split over the publicly available RosBREED Crop Reference Set and three privately held RosBREED Breeding Pedigree Sets. Except for the Breeding Pedigree Sets, curated pedigree information is given for each individual. For each individual, the type of material (selected vs. unselected), the location of sampling, and quality of the results of the sample are given. For unselected seedlings, the family to which they belong is also given. For the Breeding Pedigree Sets, this information is summarized per full-sib family.

**Table S3**: Sweet cherry germplasm genotyped and used for curation workflow. All individuals are part of the publicly available RosBREED Crop Reference Set. For each individual, curated pedigree information, the type of material (selected vs. unselected), the location of sampling, and quality of the results of the sample are given. For unselected seedlings, the family to which they belong is also given.

**Table S4**: Parameter settings used for (A) filtering SNPs used in analyses of B-allele frequency, (B) running ASSiST, (C) running FlexQTL™ for detecting Mendelian-inconsistent errors and Mendelian-consistent errors, and (D) running FlexQTL™ for phasing, haploblock determination, and creating PediHaplotyper input files.

**Table S5**: Final genetic map used for apple during data curation. For each marker, genetic position, associated haploblock, and physical position based on the apple GDDH 13 v1.1 genome are given.

**Table S6**: Final genetic map used for peach during data curation. For each marker, genetic position, associated haploblock, and physical position based on the peach v2 genome are given.

**Table S7**: Final genetic map used for sweet cherry during data curation. For each marker, genetic position, associated haploblock, and physical position based on the peach v2 genome are given.

**Table S8**: SNP classification for apple. Each SNP is classified as follows: Type ‘1’ for SNPs with good clustering and less than 5% call errors, ‘2’ for SNPs with shifted clusters causing one of the clusters to be called incorrectly, ‘3’ for SNPs with additional clusters (excluding null-alleles) that cause the incorrect identification of at least one cluster, ‘4’ for SNPs with null-alleles that cannot be correctly called automatically, ‘5’ for SNPs that could not be mapped accurately but had correct clustering, ‘6’ for monomorphic SNPs, and ‘7’ for failed SNPs.

**Table S9**: SNP classification for peach. Each SNP is classified as follows: Type ‘1’ for SNPs with good clustering and less than 5% call errors, ‘2’ for SNPs with shifted clusters causing one of the clusters to be called incorrectly, ‘3’ for SNPs with additional clusters (excluding null-alleles) that cause the incorrect identification of at least one cluster, ‘4’ for SNPs with null-alleles that cannot be correctly called automatically, ‘5’ for SNPs that could not be mapped accurately but had correct clustering, ‘6’ for monomorphic SNPs, and ‘7’ for failed SNPs.

**Table S10**: SNP classification for sweet cherry. Each SNP is classified as follows: Type ‘1’ for SNPs with good clustering and less than 5% call errors, ‘2’ for SNPs with shifted clusters causing one of the clusters to be called incorrectly, ‘3’ for SNPs with additional clusters (excluding null-alleles) that cause the incorrect identification of at least one cluster, ‘4’ for SNPs with null-alleles that cannot be correctly called automatically, ‘5’ for SNPs that could not be mapped accurately but had correct clustering, ‘6’ for monomorphic SNPs, and ‘7’ for failed SNPs.

**Figure S1**: SNP B-allele frequences plotted against physical position in the genome for (A) triploid individuals excluding ‘Jonagold’, and (B) individuals with a tetraploid pattern

**Figure S2**: Call rates observed for individuals classified as having good, intermediate, or bad quality of genotypic data as defined by their B-allele frequency plot outcome. Higher call rates are observed for individuals with better quality of genotypic data.

**Document S1**: R-script used to create B-allele frequency plots for all genotyped individuals.

**Document S2**: R-scripts used to confirm and deduce P(P)C relationships.

**Document S3**: Hands-on guideline on how to perform data curation using the steps described in this study

